# Inherited DNA damage generates multi-allelic mutations in *C. elegans*

**DOI:** 10.64898/2026.08.26.747097

**Authors:** Thomas A. Sasani, Aaron R. Quinlan

## Abstract

Exogenous and endogenous mutagens generate a wide variety of DNA lesions, including bulky adducts, chemical modifications, and single- or double-stranded breaks. A phenomenon called “lesion segregation,” in which lesions evade repair and persist for multiple cell divisions, has recently been documented in tumors and healthy somatic tissues from mice and humans, respectively. Persistent lesions can generate multi-allelic variants (MAVs) by serving as templates for multiple rounds of error-prone replication. By reanalyzing data from a large *C. elegans* mutagenesis experiment, we observed robust evidence for MAVs at a small fraction (∼0.2%) of mutated sites in the offspring of strains treated with alkylating agents. Because these sequencing data were derived from the progeny of a single F1 animal — itself the offspring of a mutagenized P0 — all mutations should be biallelic. The presence of multi-allelic variation implies that some DNA lesions are transmitted to the F1 zygote, evade repair, and are repeatedly bypassed by error-prone polymerases during embryogenesis. We suspect that many more lesions are inherited than is suggested by MAV prevalence, and that a large fraction of biallelic mutations are also caused by inherited lesions. Our results demonstrate that DNA lesions serve as durable, transgenerational templates for mutagenesis in *C. elegans*. We speculate that lesion segregation in the early embryo may be a source of mosaicism and genetic diversity in humans, as well.

## Introduction

Myriad sources of damage leave behind lesions on DNA molecules. While many lesions are efficiently repaired, others survive long enough to be encountered by DNA polymerases during cell division. These high-fidelity polymerases are often unable to replicate across lesions, triggering the recruitment of specialized translesion synthesis (TLS) polymerases [1–3]. Owing to their unique structural features, including large, accommodating active sites and a lack of exonuclease proofreading abilities, TLS polymerases are capable of synthesizing over DNA adducts, abasic sites, and other lesions that might normally lead to polymerase stalling and replication fork collapse [4–8]. However, the permissive nature of translesion synthesis comes at a cost. While many TLS polymerases are highly specialized and replicate across their cognate lesions with relatively high fidelity, TLS is more error-prone than synthesis across undamaged DNA [2,3,9,10]. TLS polymerases can also introduce new mutations downstream of lesions in a process called “collateral mutagenesis” [11–13].

### Lesion segregation and multi-allelic mutations

Aitken *et al.* (2020) [14] previously documented a phenomenon called “lesion segregation,” in which DNA lesions persist unrepaired for at least one cell division following a mutagenic insult. As long as a lesion evades DNA repair machinery, it can serve as a template for replicative enzymes, including error-prone translesion synthesis polymerases. In one round of DNA replication, an incorrect base might be incorporated opposite a lesion; in a subsequent round of replication, another incorrect base might be incorporated (Figure 1). Persistent lesions therefore become “engines of genetic diversity” by generating multi-allelic mutations [13,14]. Remarkably, some lesions can persist for years in human blood, liver and bronchial tissue [15].

**Figure 1:**
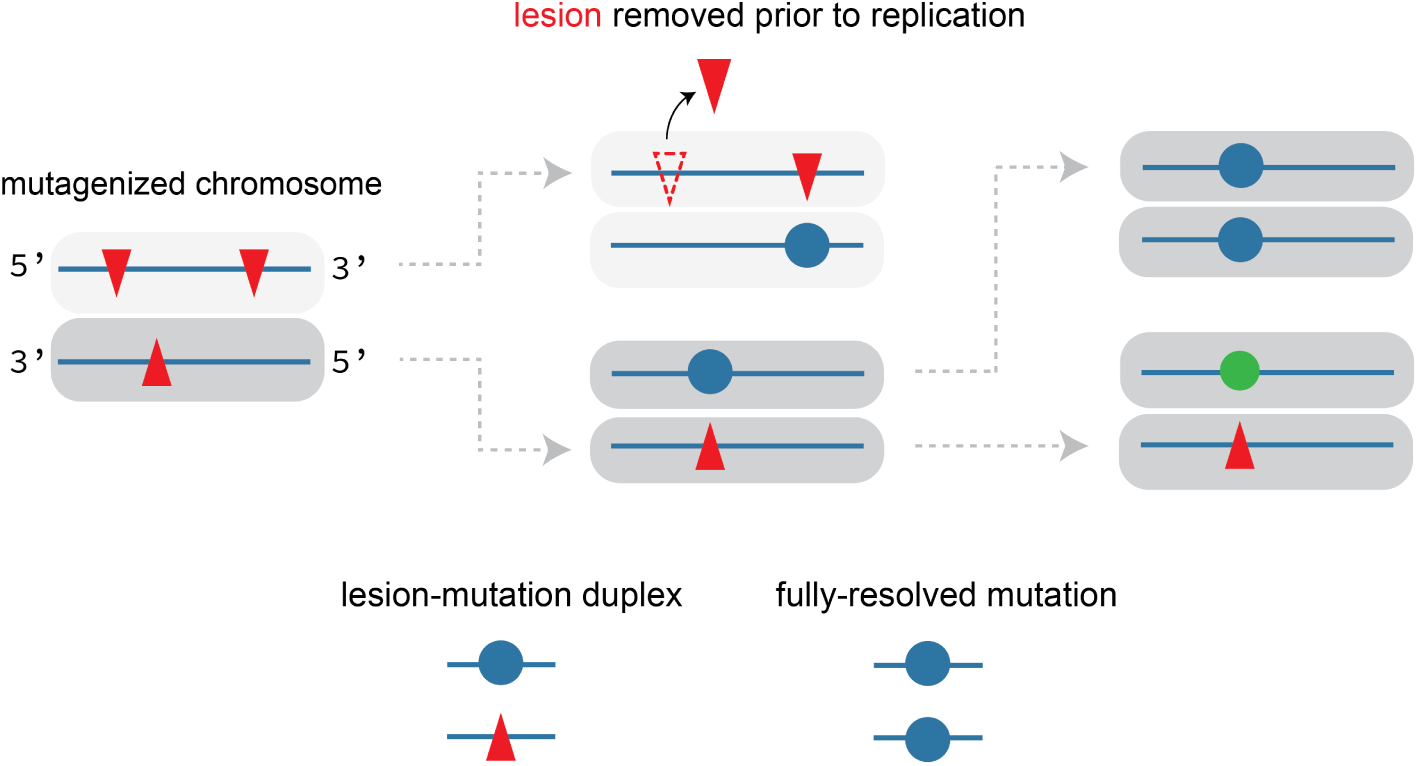
Persistent lesions can create multi-allelic mutations. DNA lesions (shown as red triangles) occur after exposure to an exogenous or endogenous mutagen. If lesions are not removed or repaired prior to DNA replication, error-prone translesion synthesis (TLS) polymerases might incorporate an incorrect base opposite the lesion, creating a lesion-mutation duplex. Lesion-mutation duplexes resolve into double-stranded mutations following a subsequent round of DNA replication. If a lesion continues to evade DNA repair machinery, it may eventually serve as a template for the incorporation of a different incorrect base (shown as a green circle). Figure adapted from [13] and [14].

### A large-scale mutagenesis experiment in *C. elegans*

Previously, Volkova *et al.* [16] mutagenized 54 *Caenorhabditis elegans* strains with 12 DNA damaging agents. Most of these strains harbored homozygous loss-of-function alleles in genes related to DNA replication and repair, including translesion synthesis (e.g., *polk-1*) and nucleotide excision repair (e.g., *xpc-1*).

Here, we reanalyze sequencing data from these mutagenesis experiments and find compelling evidence for multi-allelic variants (MAVs) in the offspring of many strains. We are confident that these MAVs are not due to sequencing errors or alignment artifacts. Instead, we believe that MAVs arise when a lesion occurs in a parental (P0) gamete, is transmitted to an F1 zygote, and persists long enough during embryogenesis to serve as a template for the incorporation of multiple *de novo* alleles. Given the invariant developmental lineage of *C. elegans* [17,18], we estimate that an inherited lesion must persist for at least five embryonic F1 cell divisions in order to generate multi-allelism in the gametes of that F1. However, lesions need only segregate unrepaired for a single cell division to generate biallelic *de novo* mutations in the F1 germline. Certain mutagens appear to cause greater levels of multi-allelism than others, though the stochastic nature of post-zygotic lesion segregation and the effects of survivor bias make it difficult to confidently interpret these differences. Nevertheless, our results suggest that DNA damage can persist from generation to generation in *C. elegans*. These persistent lesions become potent engines of mutagenesis, influencing the genetic diversity of future generations long after they arise.

## Results

### Re-analyzing a *C. elegans* mutagenesis experiment

In Volkova *et al.* (2020), 54 strains of *Caenorhabditis elegans* nematodes were either a) maintained for between 5 and 40 generations in mutation accumulation (M.A.) experiments or b) treated with various concentrations of DNA damaging agents.

#### Box 1

**Mutagenesis strategy in Volkova et al. (2020).**

P0 hermaphrodites were treated with twelve genotoxins, including alkylating agents, gamma radiation, and UV-B. In most experiments, L4 or young adult (YA) worms were treated with the genotoxin of interest, allowed to recover for 24 hours, and transferred to fresh plates. After 6 hours of egg-laying, P0 adults were removed. Two F1 animals at the L4 stage (the offspring of the mutagenized P0s) were transferred to individual plates and allowed to proliferate. A single one of these F1 lines was then expanded and used for DNA extraction and sequencing. Typically, three replicate lines were created for each genotoxin/genotype experiment.

Inspired by recent work [13–15], we hypothesized that a potent burst of genotoxins could create DNA lesions that persist for multiple cell divisions in *C. elegans*. Because a single F1 – the “child” of a mutagenized P0 – was used to initiate the population of worms used for sequencing (see Box 1), we should only observe mutations derived from lesions that were present in a single P0 sperm or egg cell (or the progenitors of those cells). As it’s succinctly described in Volkova *et al.* (2020):

> *“The zygotes which lead to the F1 generation provide a single cell bottleneck where mutations of exposed male and female germ cells are fixed before being clonally amplified during C. elegans development and passed on to the next generation in a Mendelian ratio.”*

After genotoxin treatment and sequencing, Volkova *et al.* (2020) performed read alignment and called single-nucleotide, insertion/deletion (indel), and structural variants (SVs). In total, the authors identified 135,348 high-confidence *de novo* single-nucleotide variants (SNVs) in 2,717 strains treated with genotoxins or maintained in M.A. experiments (Figure 2). Here, we limit our analyses to the subset of SNVs observed in strains treated with three alkylating agents: dimethyl sulfate (DMS), ethyl methanesulfonate (EMS), and methyl methanesulfonate (MMS). We chose these genotoxins for three reasons: first, because they primarily generated single-nucleotide and multi-nucleotide (as opposed to indel or SV) mutations; second, because they were among the most potent mutagens used in the study; and third, because worms treated with these three genotoxins were mutagenized at the same life stage (young adult; YA). We further filtered the SNVs in DMS, EMS, and MMS-treated strains to remove variants in annotated low-complexity sequences (**Materials and Methods**), producing a final callset of 68,796 alkylation-induced SNVs (Figure 2). We expect these SNVs to be biallelic, as each sequenced population of worms was derived from a single F1 animal.

**Figure 2:**
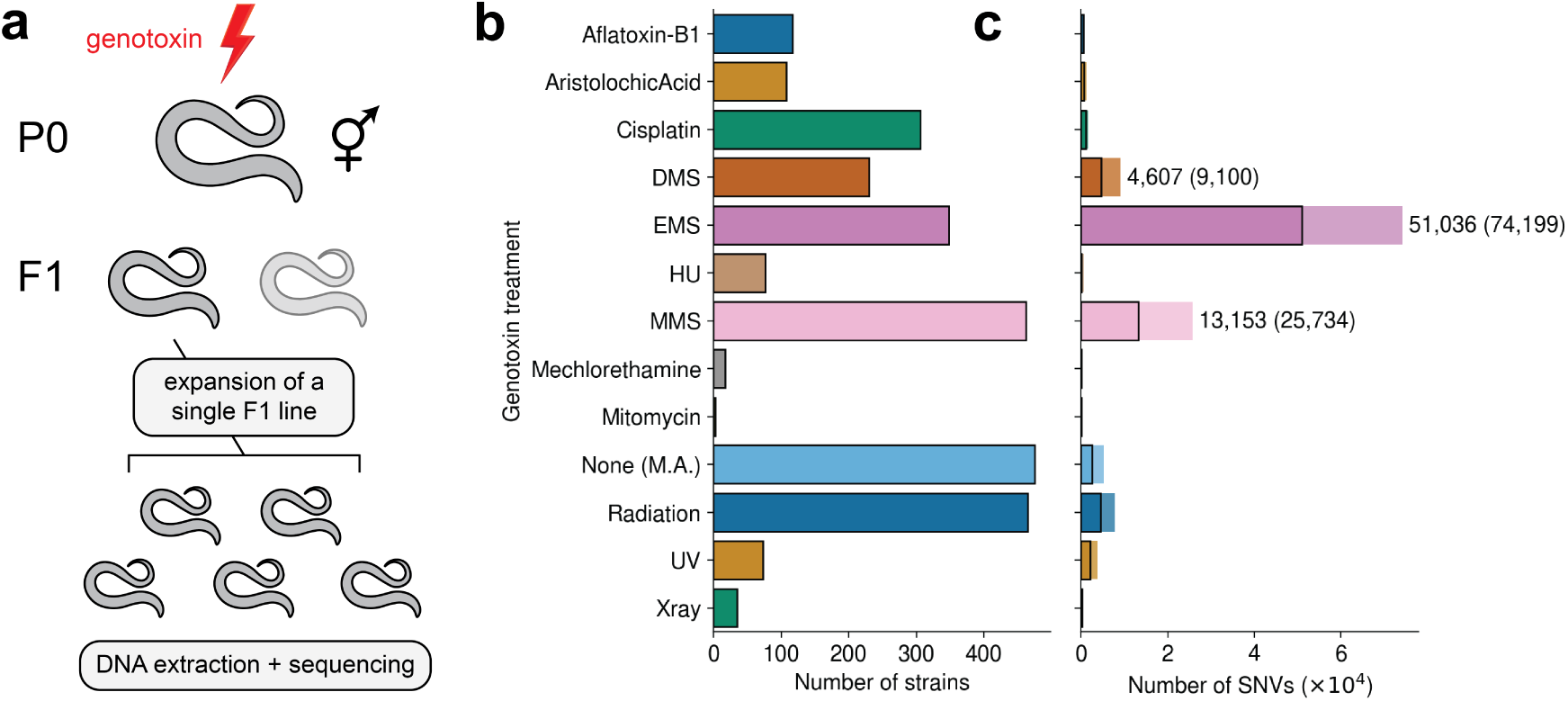
Number of genotoxin-treated strains and aggregate mutation counts in Volkova *et al.* (2020). **a)** Simplified schematic of mutagenesis strategy in Volkova *et al.* (2020). P0 worms were treated with the desired dose of genotoxin and allowed to recover for 24 hours. After 6 hours of egg-laying, adult P0s were removed. Two F1s were singled onto individual plates and allowed to proliferate. The expanded progeny of a single F1 line were used for DNA extraction and sequencing. **b)** Number of strains corresponding to each mutagenesis or mutation accumulation (M.A.) approach. DMS: dimethyl sulfate, EMS: ethyl methanesulfonate, HU: hydroxyurea, MMS: methyl methanesulfonate. **c)** Number of SNVs attributed to each mutagen, aggregated across all mutagenized strains. Darker bars (outlined in black) indicate numbers of filtered SNVs, while lighter bars indicate the total number of SNVs before stringent filtering (see **Materials and Methods**). Numbers to the right of each bar indicate the number of filtered SNVs used in this study (unfiltered counts are shown in parentheses). In this study, we exclusively analyzed mutations caused by the alkylating agents DMS, EMS, and MMS.

### Multi-allelism at mutated sites

By inspecting aligned sequencing reads at mutated sites in the genotoxin-treated lines, we observed evidence for numerous multi-allelic variants (MAVs) (Figure 3). We found almost a thousand ostensibly biallelic SNVs with at least one read supporting a “third allele,” though the number of candidate MAVs declined substantially if we required additional reads as evidence (Figure 3). To obtain a high-confidence MAV callset, we required MAVs to be supported by at least 3 sequencing reads, leaving 139 MAVs out of 68,796 total SNVs (∼0.2%).

**Figure 3:**
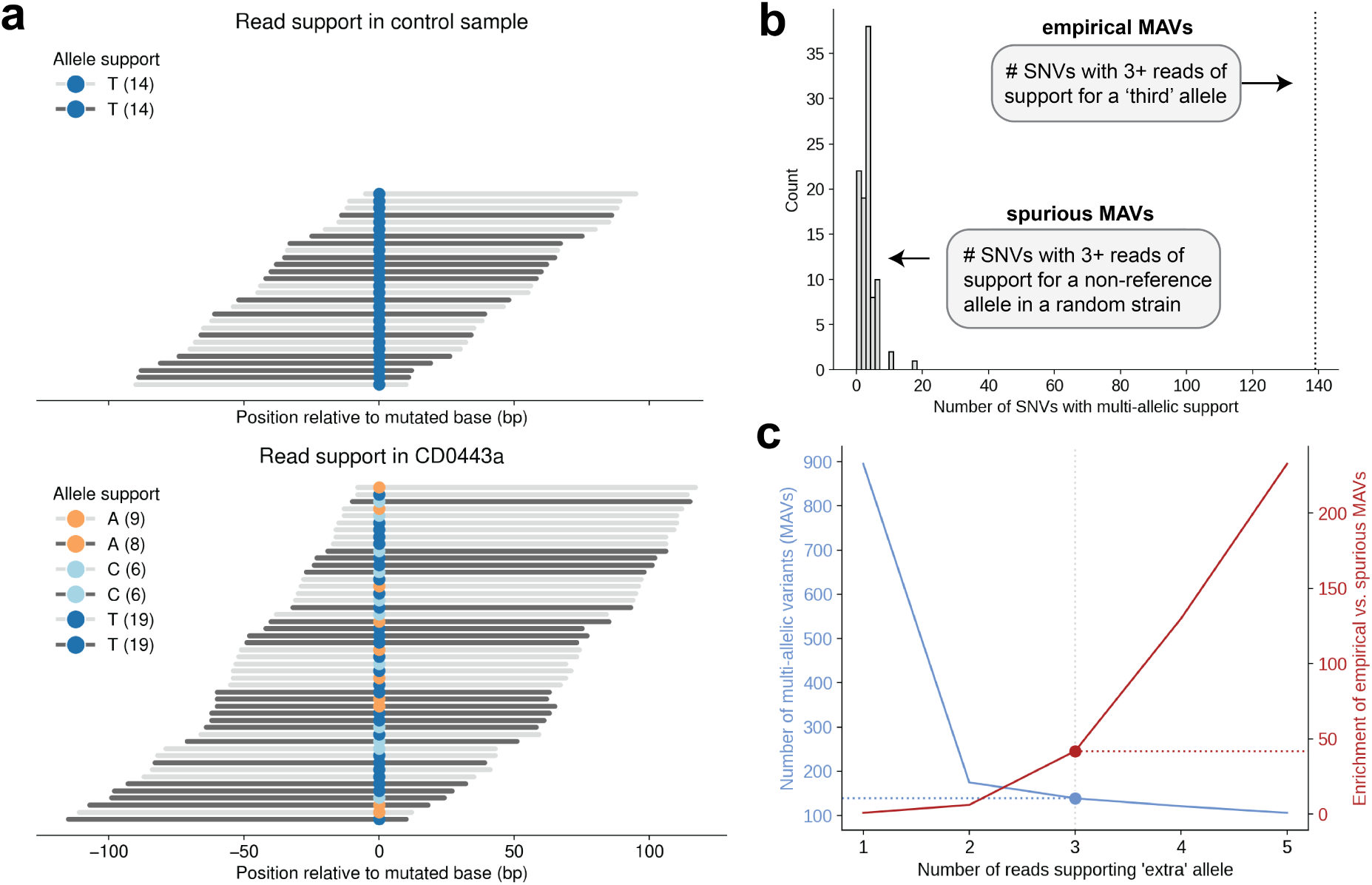
Evidence for multi-allelism in the offspring of mutagenized *C. elegans* strains. **a)** Diagram of aligned Illumina sequencing reads at a single-nucleotide variant observed in `CD0443a`, a line derived from an *xpf-1* mutant strain treated with 0.4mM MMS, and an untreated control strain. Sequencing reads are sorted by start position and colored by read orientation (light grey: aligned to the forward strand, dark grey: aligned to the reverse strand). Bases aligned to the mutated reference nucleotide are colored. For simplicity of visualization, we show a maximum of 50 reads in each subplot. Legend shows the count of each allele aligned to the forward or reverse strand. **b)** Dotted vertical line shows the number of SNVs at which we observed >=3 reads supporting a third allele. In each of 100 trials, we calculated the number of SNVs at which we observed evidence for multi-allelism (>= 3 reads supporting a non-reference allele) using a random strain’s sequencing reads. The distribution of “spurious” MAVs identified in each trial is shown as a grey histogram. **c)** We performed the random sampling procedure described in **b)** using varying thresholds on the number of reads required to support a candidate MAV. The blue line shows the number of empirical MAVs identified at each read threshold. The red line shows the enrichment of empirical vs. spurious MAVs identified at each threshold, calculated by dividing the total number of empirical MAVs by the average number of spurious MAVs identified across 100 random sampling trials.

The Volkova *et al.* (2020) strains were sequenced with two different Illumina platforms (HiSeq 2000 and HiSeq X Ten). Nearly all of the strains processed on the HiSeq X Ten were sequenced to an average of ∼30X depth, while those processed on HiSeq 2000 were sequenced to an average of ∼90X (Supplementary Figure 1). To determine whether sequencing coverage significantly affected our ability to detect MAVs, we randomly downsampled every strain to an average of ∼30X (**Materials and Methods**). We detected 91 MAVs using the downsampled alignments, demonstrating that MAV detection requires relatively high sequencing depth, and suggesting the true number of MAVs in the Volkova *et al.* (2020) strains is likely greater than our estimate here.

We were initially suspicious that MAVs were simply a result of sequencing errors or alignment artifacts. In each DMS, EMS, or MMS-treated strain, we calculated the number of SNVs at which 3 or more reads supported a non-reference allele in a random strain (Box 2, **Materials and Methods**).

#### Box 2

**Generating a null expectation for the number of MAVs.**

To identify “empirical” multi-allelic variants (MAVs), we first searched for multi-allelic evidence at the set of *M* SNVs (where |*M*| = 68,796) in 691 strains treated with DMS, EMS, or MMS. In a given strain, we asked whether there were 3+ sequencing reads supporting a ‘third’ allele at each of the strain’s ostensibly biallelic SNVs *S*; *S* ∈ *M*. If so, we classified the site as an MAV.

We then calculated a null expectation for the number of MAVs we’d expect to see in a given strain due to sequencing error, alignment artifacts, and so on. For a given strain with *S* ostensibly biallelic SNVs, we randomly sampled a different strain’s read alignments (from the complete collection of *n* = 2,717 strains, excluding those with the same genotype as the focal strain, as some de novo mutations may have fixed in a strain prior to genotoxin treatment). We then used that random strain’s alignments to calculate evidence for multi-allelism at each of the *S* variants in the original strain. Since a random strain should be homozygous for the reference allele at every variant observed in the focal strain, we considered any non-reference evidence to be evidence for a “spurious” MAV. We repeated this process (randomly sampling a different strain’s alignments and re-calculating evidence for spurious MAVs) 100 times.

Compared to this null expectation, MAVs occurred far more often than expected by chance (Figure 3b). We observed an average of 3.3 spurious MAVs per random sampling trial, and never observed more than 18 spurious MAVs in a single trial. Even after removing SNVs in annotated low-complexity regions (**Materials and Methods**), we found that spurious MAVs occurred in lower-complexity sequences than “empirical” MAVs (two-sided Kolmogorov-Smirnov *p* = 3.0 × 10^−3^; Supp. Fig. 2). There was no difference in sequence complexity between empirical MAVs and biallelic SNVs (two-sided KS *p* = 0.29; Supp. Fig. 2). Because many of the strains in Volkova *et al.* (2020) were barcoded and sequenced in multiplexed Illumina runs, we also searched for evidence that “index hopping” (a phenomenon in which reads originating from one sample are incorrectly assigned to another sample during sequencing [19]) could explain the presence of MAVs (**Materials and Methods**). We found no evidence that the “extra” alleles at MAVs were present in other strains, indicating that index hopping was unlikely to explain multi-allelism at SNVs. Finally, we observed similar levels of strand bias at empirical MAVs and biallelic SNVs (*χ*^2^ *p* = 0.37; **Materials and Methods**).

### All tested alkylating agents create multi-allelism

Using our stringently filtered callset, we measured the degree to which different genotoxins generated multi-allelic variants. Compared to the null expectation, MMS caused the greatest enrichment of MAVs in *C. elegans* (approximately 0.7% of all MMS-induced SNVs were multi-allelic) (Figure 4). A substantial fraction of DMS-induced SNVs were multi-allelic, as well (∼0.5%). Despite generating the largest number of SNVs overall, EMS generated relatively few MAVs (∼0.05%). It is challenging to attribute these differences in MAV enrichment to the characteristics of specific mutagens, however (see **Discussion**).

**Figure 4:**
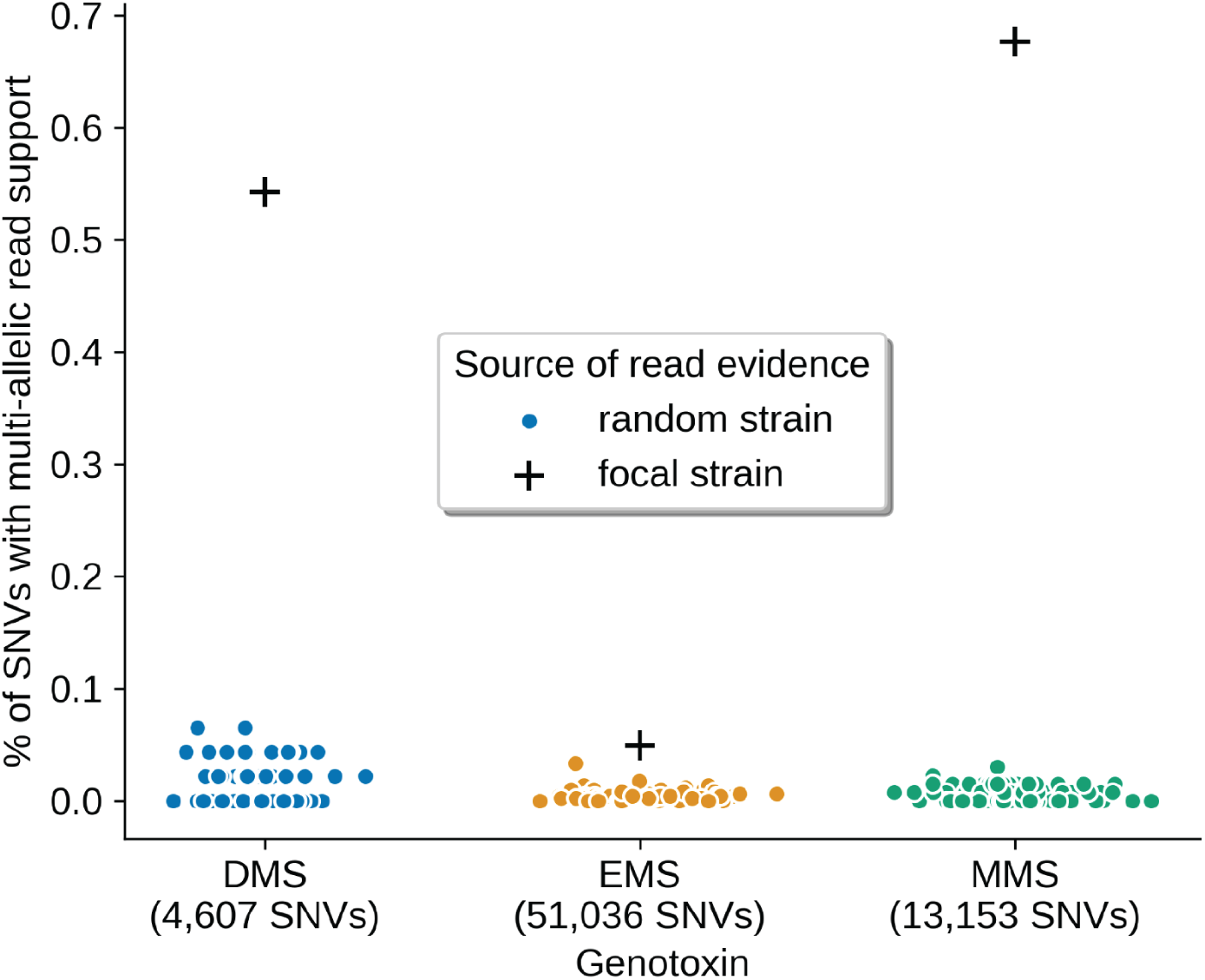
Enrichment of MAVs after treatment with three alkylating agents. Using *n* = 691 strains treated with DMS, EMS, or MMS, we calculated the proportion of SNVs at which we observed 3 or more reads supporting a multi-allelic variant (MAV). The total number of empirical MAVs we identified in strains treated with a given mutagen is shown as a black “plus” sign. We then performed 100 random sampling trials to obtain a null expectation for the number of observed MAVs (see Box 2). In each trial, we calculated the proportion of spurious MAVs identified across strains treated with the specified mutagen. The distribution of these spurious MAV proportions across 100 trials is shown using a colored jitterplot.

We provide a table of all observed MAVs observed in **Supplementary Table 1**, and have deposited interactive, self-contained Integrative Genomics Viewer HTML reports [20] for each MAV on GitHub.

### A simple model of transgenerational DNA damage

Convinced that the observed multi-allelic mutations were real sequence variants, we next considered how they might occur (Figure 5). Let’s first assume that genotoxin treatment occurs at the young adult stage (YA) in a P0 animal.

**Figure 5:**
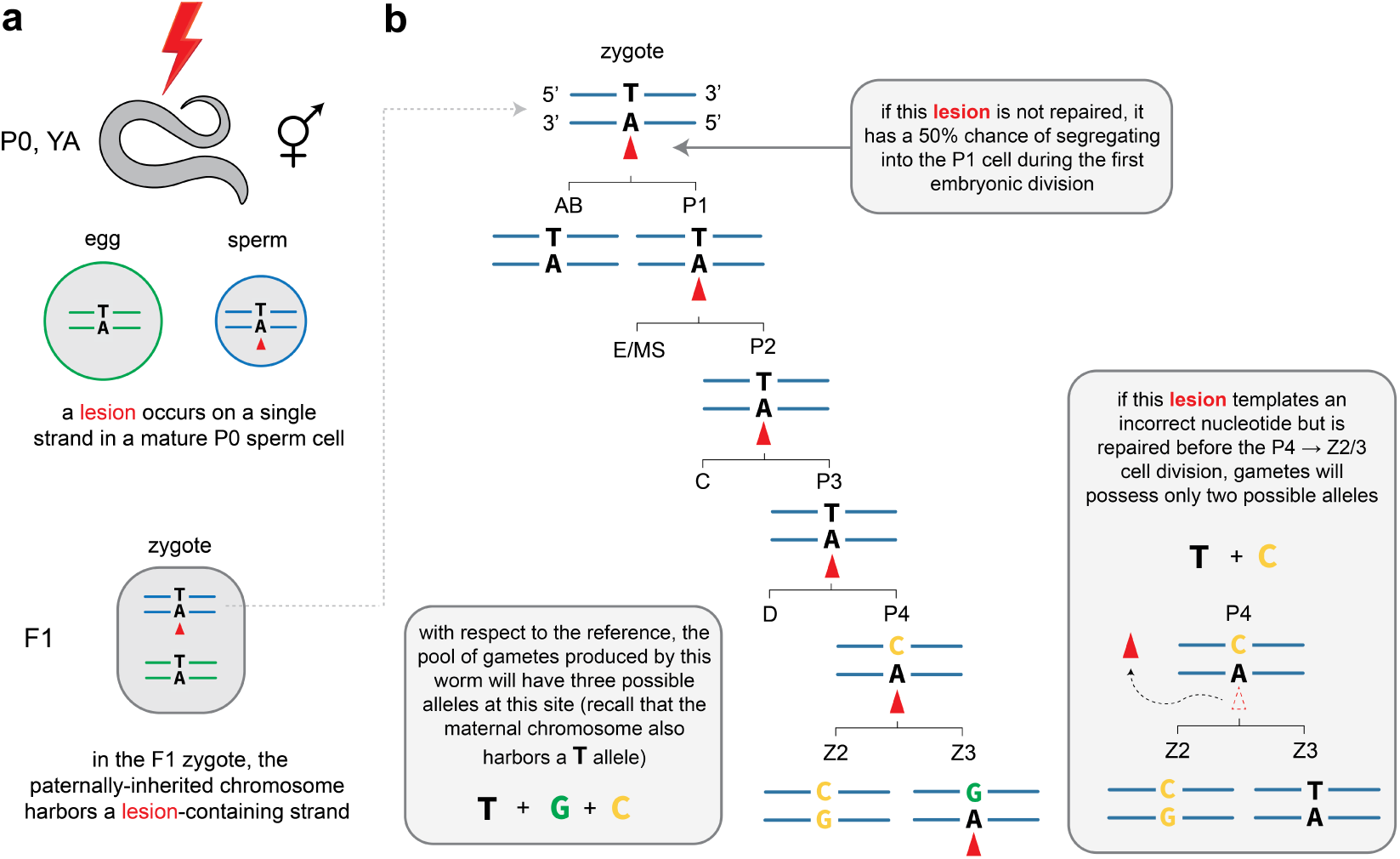
Lesions must segregate unrepaired into the *Z*_2_ or *Z*_3_ cell to generate multi-allelism. In this toy diagram, we track the fate of a single lesion-containing DNA strand during the first few cell divisions of *C. elegans* embryogenesis. **a)** Following genotoxin treatment in a young adult (YA) worm, a lesion (shown as a red triangle) is generated in a haploid gamete. Here, a lesion is created at an adenine on the reverse strand of a haploid sperm cell’s genome. A damage-free egg cell is then fertilized by the sperm cell that harbors a DNA lesion. **b)** The sperm-derived lesion could persist during the first few F1 embryonic cell divisions. If it evades repair, the lesion will segregate into either the *AB* or *P*_1_ cell (the two products of the first embryonic cell division). If the lesion-containing strand continues to segregate into germline progenitor cells (*P*_2_, *P*_3_, and *P*_4_) without being repaired, it could template the incorporation of multiple unique derived alleles during gametogenesis. Here, the lesion templates the incorporation of a **`C`** allele during the *P*_3_ → *P*_4_ division, and a **`G`** allele during the 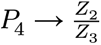 division. If the lesion templates the incorporation of an incorrect base but either a) segregates into the somatic lineage or b) undergoes error-free repair prior to the 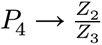 cell division, the F1′s gametes could be mosaic for two alleles, and the mutation would appear biallelic in future generations (see inset). In this toy example, we only depict the fate of the chromosome donated by the parental sperm cell. Recall that each cell also possesses a copy of the maternal chromosome — with a T:A base pair at the focal site — that contributes to the eventual population of F1 gametes.

Let’s next imagine that in the mutagenized P0, a mature, haploid sperm cell harbors a DNA lesion (Figure 5). After that sperm cell fertilizes an egg, what are the chances its lesion would persist long enough to create multi-allelism in the F1′s gametes? To answer this question, we can take advantage of the fact that the *C. elegans* embryonic lineage is essentially invariant [17,18]. All *C. elegans* gametes are derived from a single germline progenitor cell called *P*_4_. The *P*_4_ cell divides to create *Z*_2_ and *Z*_3_, which localize to the gonad and give rise to all eventual sperm and oocytes [21]. If the lesion segregates into the somatic lineage or undergoes error-free repair prior to the 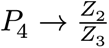 cell division — and never templates the incorporation of an incorrect base — the F1 gametes will only possess a single allele at that site, and any evidence of an inherited lesion will be lost (Supp. Fig. 3a). But what happens if an incorrect base *is* incorporated opposite the lesion?

For example, let’s consider the simplest possible way for an inherited lesion to generate multi-allelism in the next generation. First, imagine that the lesion undergoes error-prone translesion synthesis (TLS) and a **`C`** is incorporated opposite the lesion during the *P*_3_ → *P*_4_ division. If the lesion is again bypassed by error-prone TLS during the 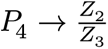 division, both the first and second incorrect bases ( **`C`** and **`G`**) will be present in mature gametes. For the sake of example, we assume that error-prone TLS is the mechanism by which mutations are introduced opposite lesions. One could also imagine other ways for *de novo* mutations to occur, though these processes might result in the lesion being removed. For example, during the 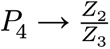 division, nucleotide excision repair machinery might remove the lesion and fill the gap with an incorrect nucleotide. Though the lesion would disappear, both *de novo* alleles would propogate into mature gametes.

It’s also possible for an inherited lesion to generate mutations that appear *biallelic* in subsequent generations. For example, if a lesion templates a **`C`** during the *P*_3_ → *P*_4_ division, but is repaired prior to 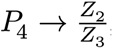, only the **`T`** and **`C`** will exist in the F1′s mature gametes. We believe that this simple model, in which a lesion-containing strand persists for multiple embryonic cell divisions, explains the presence of MAVs in the Volkova *et al.* (2020) dataset.

## Discussion

In this analysis, we provide evidence that genotoxin treatment creates DNA lesions that persist from generation to generation in *C. elegans* nematodes. In line with recent results [22], we find that DNA damage in a P0 animal can produce a measurable transgenerational phenotype (here, the presence of multi-allelic variation in the progeny of an F1). Our work suggests that inherited lesions serve as durable, persistent templates for *de novo* mutagenesis in *C. elegans*.

### Prior evidence for the transgenerational effects of DNA damage in *C. elegans*

Wang et al. [22] recently demonstrated that paternal exposure to ionizing radiation leads to transgenerational embryonic lethality in *C. elegans*. In their study, irradiated male worms were mated to healthy, “feminized” hermaphrodites. While the F1 offspring of these irradiated males were mostly viable, a large proportion of F2s exhibited embryonic lethality. The authors concluded that DNA damage — primarily in the form of double-stranded breaks (DSBs) — was transmitted from P0 sperm cells to the zygotes of the F1 generation. In the F1 germline, these damaged chromosomes triggered the formation of heterochromatin, preventing error-free mechanisms (like homologous recombination) from repairing DSBs and leading to embryonic lethality in F2s [22]. These results demonstrated that radiation-induced DNA damage persists from parental gametes to zygotes, and that inherited DNA damage can cause epigenetic changes that significantly impact the viability of future generations. Here, we describe a related — but distinct — phenomenon, in which single-nucleotide DNA lesions are themselves inherited from generation to generation, serving as recurrent templates for *de novo* mutations and generating heritable genetic diversity.

### How likely is gametic mosaicism due to inherited lesions?

The probability that a DNA lesion occurs on both strands at the same nucleotide in a P0 gamete (or, that it occurs at the same nucleotide in both gametes) is extremely small. Thus, we can safely assume that a given MAV is caused by the presence of a single lesion on a single DNA strand in a zygote. We can also estimate the probability that a lesion-containing strand would segregate into a *P*_4_ cell. Assuming the lesion evades DNA repair machinery during each cell cycle, there is a 50% chance the lesion-containing strand will be inherited from a zygote to the *P*_1_ cell, a 50% chance it will be inherited from *P*_1_ to *P*_2_, and so on (Figure 5). Thus, the probability that a lesion is inherited by the *P*_4_ cell (assuming it is not repaired) is simply 0.5^4^ = 0.0625. Lesion segregation in the early zygote is therefore stochastic, depending on the vagaries of Mendelian segregation.

### Missing MAVs due to the duration of lesion segregation and low sequencing depth

If it segregates into a *Z*_2_ or *Z*_3_ cell and continues to evade repair, a lesion could also create multi-allelic variants by templating incorrect nucleotides during the later stages of germ cell development (Supp. Fig. 3b). These MAVs would be present in correspondingly fewer mature gametes, and therefore present in a smaller fraction of the F1′s progeny. If we were to sequence a small number of pooled F2s, or sequence those pooled F2s to relatively low depth, the MAV might escape detection. Indeed, we detected only 91 of 139 MAVs after downsampling the Volkova *et al.* (2020) sequencing data to 30X (**Materials and Methods**).

Many lesions likely create *de novo* mutations that appear biallelic in subsequent generations. For example, if a lesion is incorrectly bypassed during the *P*_3_ → *P*_4_cell division, a lesion-mutation duplex would be present in the *P*_4_ cell (see inset in Figure 5). But if the lesion is repaired during the next cell cycle, the gametes produced by that worm would only contain two possible alleles, and its offspring would appear to be biallelic (Figure 5). Similarly, an incorrect base could be incorporated opposite the lesion during the zygote → *P*_1_ (or *P*_1_ → *P*_2_, or *P*_2_ → *P*_3_) division. If the strand containing the incorrect base continued to segregate into the *P* lineage — while the lesion-containing strand segregated into a somatic progenitor — the resulting mutation would be fully-resolved (i.e., the incorrect nucleotide and its complement would be present on both strands) in the *P*_4_ cell (Supp. Fig. 3c). Exactly half of the F1′s gametes would possess the incorrect base, and the site would appear to be biallelic among the F2s. Thus, we suspect that a substantial fraction of biallelic SNVs observed in the Volkova *et al.* (2020) dataset likely result from inherited lesions. The early zygote probably inherits far more damage than is suggested by MAV prevalence.

### Missing MAVs due to survivor bias

MAVs might also be invisible due to the strength of genotoxin treatment. If a P0 animal were treated with an extremely high dose of mutagen, its F2 progeny could exhibit very high embryonic lethality as a consequence of newly homozygous loss-of-function alleles. If an F1′s gametes are mosaic for multiple derived alleles at a given site, fewer F2 progeny provide us with fewer opportunities to observe those “extra” alleles. In other words, MAV detection is complicated by survivor bias; high mutagen doses create more lesions, and these lesions might generate multi-allelic variation, but those multiple alleles may not segregate into living offspring. Ultimately, then, our ability to detect MAVs depends on at least four distinct factors: 1) the random segregation of lesion-containing strands into germline progenitors, 2) the efficiency of lesion repair in the early embryo (i.e., the duration of lesion persistence), 3) the fidelity of translesion synthesis, and 4) the reproductive success of the F1 and its progeny following mutagen exposure.

### What mechanisms likely lead to multi-allelism?

As suggested in previous studies of lesion segregation [13,14], we hypothesize that multi-allelic variants (MAVs) arise from error-prone replication over persistent lesions. In the span of about two hours, the newly formed *C. elegans* zygote undergoes five asynchronous and asymmetrical cell divisions [23]. The early embryo is highly sensitive to the timing of these divisions, and cell cycle delays can lead to embryonic lethality [23]. *C. elegans* embryos therefore rely on translesion synthesis (TLS) polymerases to bypass DNA damage that might otherwise stall replication forks and activate cell cycle checkpoints [24–26]. As a result, inherited lesions may be able to persist for multiple embryonic cell divisions, and each division presents an opportunity for a new, incorrect base to be incorporated. We hypothesize that a reliance on TLS for DNA damage bypass makes the early *C. elegans* embryo susceptible to lesion segregation and the generation of heritable MAVs.

### Do some mutagens create more persistent lesions?

Compared to a null expectation, we found that certain mutagens led to a greater enrichment of MAVs than others. For example, despite causing the highest number of mutations (Figure 2), EMS treatment led to a relatively modest enrichment of MAVs. Without additional experimental data, it is difficult to conclusively say whether genotoxins like DMS and MMS are more potent engines of multi-allelism or create longer-lasting DNA lesions. As discussed above, our ability to detect MAVs relies on a combination of factors: the chance segregation of lesion-containing strands into germline progenitors, the efficiency of embryonic DNA repair, and the lethality induced by a mutagen itself. Mutagens like EMS almost exclusively create C:G → T:A mutations, and often generate premature stop codons, strong loss-of-function variants, and null alleles [27,28]. EMS mutagenesis may therefore prove more lethal to F2 embryos, limiting our ability to sequence the full complement of *de novo* alleles present in the offspring of an EMS-treated P0. While we likely failed to observe many of these large-effect alleles due to survivor bias (any worms with lethal null alleles were not sequenced), we did find that EMS-derived SNVs occurred at more conserved nucleotides in the *C. elegans* genome (Supp. Fig. 4).

Moreover, our analysis in this manuscript deals only with single-nucleotide mutations; the mutagens we analyzed also create small numbers of insertions, deletions, and structural variants [16]. Certain mutagens are more likely to generate null alleles via these types of mutations, potentially leading to even greater F2 lethality and a reduced ability to detect MAVs.

### Implications for sex-biased mutagenesis

The Volkova *et al.* (2020) dataset is extremely powerful, though it leaves one important question untested: are lesions more likely to be inherited from sperm or egg cells? Wang et al. (2023) [22] hypothesize that the compacted chromatin architecture of *C. elegans* sperm [29] renders them incapable of repairing damage caused by ionizing radiation. In this study, we analyzed alkylating mutagens that were applied to P0 hermaphrodites at the early young adult (YA) stage of development [16]. In *C. elegans* hermaphrodites, spermatogenesis and oogenesis occur in sequence; spermatogenesis begins and ends during the L4 larval stage, while oogenesis begins afterward and continues through adulthood [21]. The P0 worms in Volkova *et al.* (2020) were allowed to recover for 24 hours after treatment with alkylating agents; following this recovery period, F1 progeny were collected within 6 hours [16]. Based on this treatment timeline, we speculate that most F1 offspring were fertilized by gametes that were either a) mature sperm or b) pachytene-stage oocytes at the time of mutagenesis [30]. MAVs probably derive from persistent lesions inherited from both sperm and oocytes, though we suspect that mature sperm are more susceptible to genotoxin treatment.

The timing of genotoxin exposure likely has a significant impact on the likelihood of lesion persistence. Wang *et al.* reported embryonic lethality in the offspring of males irradiated at the late L4 stage (when spermatogenesis is complete), but not in the offspring of males irradiated at the late L3 or early L4 stage (when spermatogenesis is ongoing) [22]. This result suggests that repair processes in a gamete progenitor can correct DNA damage relatively quickly. Future experiments, in which male and “feminized” hermaphrodites are separately mutagenized at different developmental stages, could provide insight into the relative ability of sperm and oocytes to repair DNA lesions caused by alkylating agents.

### Implications for human *de novo* mutagenesis

A recent study of human germline *de novo* mutation (DNM) reported a positive correlation between maternal age and the number of DNMs on paternal haplotypes [31]. This result has been attributed to mutations that occur during early embryogenesis. Because the early embryo relies on maternally-deposited machinery to replicate and repair DNA, older oocytes may contribute more defective machinery, leading to elevated rates of post-zygotic mutation on both maternal and paternal haplotypes [31–35]. Post-zygotic maternal repair of paternal lesions — and its role in generating a wide range of both small and structural *de novo* mutations — has been studied extensively in mice [36–38], though its contribution to human genetic diversity remains poorly characterized. We speculate that inherited DNA damage may be an underappreciated source of heritable *de novo* mutations in humans, particularly if parental gametes are exposed to a mutagen in the weeks or months prior to conception, as suggested previously [22,37].

### What about indels and SVs?

Finally, we note that multi-allelic SNVs are not the only kind of multi-allelism that might be caused by a persistent lesion. Volkova *et al.* (2020) demonstrated that genotoxins create a diverse range of DNA mutations, from SNVs and multi-nucleotide variants to insertions, deletions, tandem repeat expansions, and larger structural variants. Here, as a proof of concept, we focused on three alkylating agents that predominantly generate SNVs. Using long-read sequencing technologies, like the PacBio and Oxford Nanopore Technologies platforms, as well as new technologies that enable accurate sequencing in low-complexity regions of the genome, like the Element AVITI platform, we can begin to investigate the myriad consequences of persistent lesions on other kinds of DNA mutation. For example, a persistent lesion might template the incorporation of two unique SNV alleles, as well as a small deletion; or an SNV allele and two unique indels; and so on. We anticipate that future studies will clarify the role of persistent DNA damage in generating inherited genetic diversity.

## Acknowledgments

We thank Drs. Bettina Meier and Anton Gartner for assisting with data access. We also thank Dr. Gartner for helpful feedback on our initial findings and a draft of the manuscript. We thank members of the Quinlan Lab for helpful discussions and brainstorming. Dr. Aaron Quinlan is supported by award R01HG012252 from the National Institutes of Health (NIH). The computational resources used in this study were partially funded by the NIH Shared Instrumentation Grant 1S10OD021644-01A1.

## Materials and Methods

### Data and code availability

No new data were generated as part of this study. All sequencing data are available on the European Nucleotide Archive (study accessions `ERP000975` and `ERP004086`) and all single-nucleotide variant calls are available in VCF format in Supplementary Data File 6 from Volkova *et al.* (2020) [16]. We have distributed a Snakemake [39] pipeline that can be used to reproduce all data processing, statistical analysis, and figure generation, which can be accessed at https://github.com/tomsasani/lesion-segregation. Processed data, including filtered SNV mutation calls in each strain, filtered MAVs, and self-contained, interactive Integrative Genomics Viewer reports for each MAV [20], are also available at the above GitHub repository.

### Processing sequencing data from Volkova et al. (2020)

We downloaded FASTQ files for all relevant strains from the European Nucleotide Archive (ENA) using the fasterq-dump utility (SRA tools v3.4.1):

~~~
fasterq-dump -e 4 -p -O /path/to/fastq/dir/ -o $sample_prefix $accession
~~~

We then filtered the reads (adapter trimming, quality filtering, length filtering, etc.) with FASTP v0.20.1 [40], using the following command:

~~~
fastp --in1 /path/to/fastq.1.gz \
      --in2 /path/to/fastq.2.gz \
      --out1 /path/to/fastq.clean.1.gz \
      --out2 /path/to/fastq.clean.2.gz \
      --unpaired1 /path/to/fastq.unpaired.1.gz \
      --unpaired2 /path/to/fastq.unpaired.2.gz \
      --thread $threads \
      -V
~~~

We aligned the cleaned FASTQ files to the WBcel235/ce11 build of the *C. elegans* reference genome (downloaded from the UCSC Genome Browser) using `bwa mem` (v0.7.19-r1273) [41] and the Picard toolkit within GATK (v4.6.0) [42,43] to produce sorted, duplicate-marked BAM files:

~~~
bwa mem \
    -t $threads \
    -K 96000000 \
    -R “read_group_string” \
    /path/to/reference.fa \
    /path/to/fastq.1.clean.gz \
    /path/to/fastq.2.clean.gz \
    | \
~~~

~~~
gatk SortSam \
    --java-options -Xmx24g \
    --MAX_RECORDS_IN_RAM 5000000 \
    -I /dev/stdin \
    -O /path/to/sorted.bam \
    --SORT_ORDER coordinate \
    --TMP_DIR /path/to/tmpdir \
    --CREATE_INDEX true
~~~

~~~
gatk MarkDuplicates \
    --java-options -Xmx24g \
    -I /path/to/sorted.bam \
    -O /path/to/sorted.dupmarked.bam \
    --TMP_DIR /path/to/tmpdir \
    -M /path/to/metrics.txt
~~~

We downloaded single-nucleotide variant (SNV) VCF files for every strain (called with respect to the WBcel235/ce11 reference) from the Supplemental Material of Volkova *et al.* (2020). We performed additional filtering on these SNV calls by removing SNVs that overlapped repetitive regions annotated by WindowMasker (`windowmaskerSdust` track for `ce11` downloaded from the UCSC Table Browser). After filtering, we retained a total of 68,796 SNVs.

### Identifying multi-allelic variants

At every SNV in a particular strain, we used `pysam` v0.23.3 (https://github.com/pysam-developers/pysam) to query the strain’s BAM file at the variant position. We counted the number of sequencing reads that supported a particular single-nucleotide allele at the variant position, excluding the following:

1. reads with mapping quality < 60
2. bases with Phred-scaled base quality < 20
3. improperly paired reads
4. read pairs aligned to different chromosomes
5. supplementary alignments
6. secondary alignments
7. bases within 5bp of the aligned start or end coordinate of a read
8. reads marked as duplicates

### Calculating the expected number of multi-allelic variants

To determine whether we identified more MAVs than expected given the background rate of Illumina sequencing error and alignment artifacts, we took a simple random sampling approach. For a given “focal” strain with *S* SNV mutations, we randomly sampled a BAM file from the complete set of 2,717 strains (excluding the focal strain and any strains with the same genotype as the focal strain). We then searched for evidence of multi-allelic variants at all *S* sites using that random strain’s reads, as described above. We repeated this process 100 times for every focal strain. We considered any non-reference read evidence in a random strain as support for a “spurious” MAV. In each of the 100 trials, we calculated the total number of spurious MAVs observed across the alkylation-treated strains.

### Testing for the presence of index hopping

One possible explanation for the presence of multi-allelic variants (MAVs) is “index hopping,” a phenomenon that occurs in multiplexed Illumina sequencing runs [19]. Many of the mutagenized strains in Volkova *et al.* (2020) were barcoded and run in a multiplexed library. After sequencing, these sample barcodes were used to “demultiplex” the reads, assigning each read to the sample from whom it originated. However, during Illumina sequencing runs, sample indices can occasionally “hop” from one sample to another on a flow cell, meaning that reads from one sample are incorrectly assigned to another sample. To test for the possibility that index hopping could explain the presence of MAVs in our datasets, we iterated over each of the 139 MAVs identified in strains treated with DMS, EMS, and MMS. At each MAV, we searched for evidence of the non-reference alleles in every other strain’s alignments (n = 2,717) using `pysam` v0.23.3. We did not find any evidence that non-reference alleles at MAVs were segregating in other Volkova *et al.* (2020) strains.

### Measuring strand bias

At every biallelic SNV, we counted the number of reads supporting the alternate allele that were aligned to the forward and reverse strand. We performed a binomial test for strand bias using `scipy.stats` as follows:

~~~
binom = ss.binomtest($n_fwd, $n_fwd + $n_rev, p=0.5)
~~~

We considered a site with a binomial *p* < 0.05 to have significant strand bias. At every candidate multi-allelic variant, we performed a similar test for strand bias using the number of reads supporting each alternate allele. We then compared the count of strand-biased sites in the biallelic and multi-allelic callsets using a Chi-square contingency table:

|  | strand biased | not strand biased |
| --- | --- | --- |
| biallelic | 6,152 | 62,505 |
| multiallelic | 16 | 123 |

We observed similar numbers of biallelic and multi-allelic sites with significant strand bias (*χ*^2^ *p* = 0.37).

### Downsampling experiments

We used `mosdepth` (v0.3.14) [44] to calculate average genome-wide sequencing depth in every strain from Volkova *et al.* (2020).

~~~
mosdepth -t $threads \
         -nx \
         -f /path/to/reference.fa \
         $prefix \
         /path/to/input.cram
~~~

Most strains were sequenced to a genome-wide average of ∼90X, but many were sequenced to an average of ∼30X. Sequencing coverage was highly correlated with the platform on which samples were processed; most strains with high coverage were sequenced on the Illumina HiSeq 2000 platform, while those with lower coverage were sequenced on Illumina HiSeq X Ten. Because we require at least three reads of support to confidently detect multi-allelic variants (MAVs), increased sequencing depth would likely give us greater power to detect MAVs (especially if the “third” allele was introduced later during gametogenesis, and fewer F2 progeny harbor the allele). To test for the effects of both sequencing coverage and sequencing platform on MAV detection, we downsampled every strain’s CRAM file to an average of 30X depth using `samtools` (v1.16) [45]:

~~~
samtools view --write-index \
              -C \
              -s $downsampling_factor \
              -T /path/to/reference.fa \
              -@ $threads \
              -o /path/to/output.cram \
              /path/to/input.cram
~~~

where `$downsampling_factor` was calculated as min(0.999, (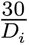)), where *D_i_* is the average genome-wide depth in strain *i*. We then searched for MAVs in each strain as described above, using downsampled CRAMs in place of the strains’ original CRAMs. We note that our downsampling experiments are imperfect, because we searched for MAVs at the ostensibly biallelic SNVs originally identified in Volkova *et al.* (2020). These SNVs were identified using the original alignments for each strain (which, in some cases, were sequenced to very high depth). For the purposes of this experiment, we assume that all of the original SNVs identified in Volkova *et al.* (2020) would be detectable using a downsampled 30X alignment.

## Supplementary Figures

**Supplementary Figure 1:**
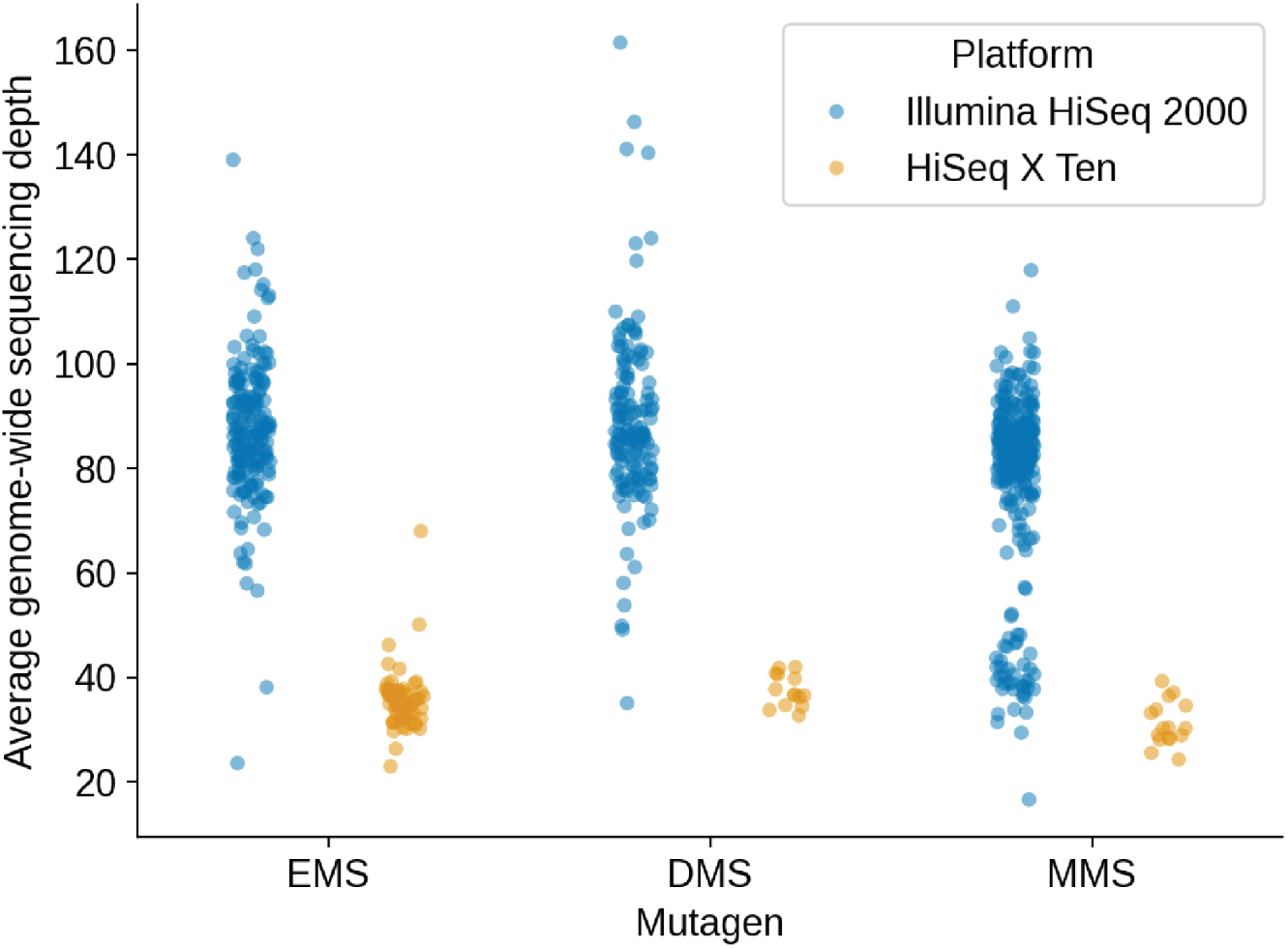
Sequencing depth across Volkova *et al.* (2020) strains. We used `mosdepth` [44] to calculate the average genome-wide sequencing depth in all strains treated with EMS, DMS, and MMS. Each point represents a single strain and is colored according to the Illumina platform (HiSeq 2000 or HiSeq X Ten) used for sequencing.

**Supplementary Figure 2:**
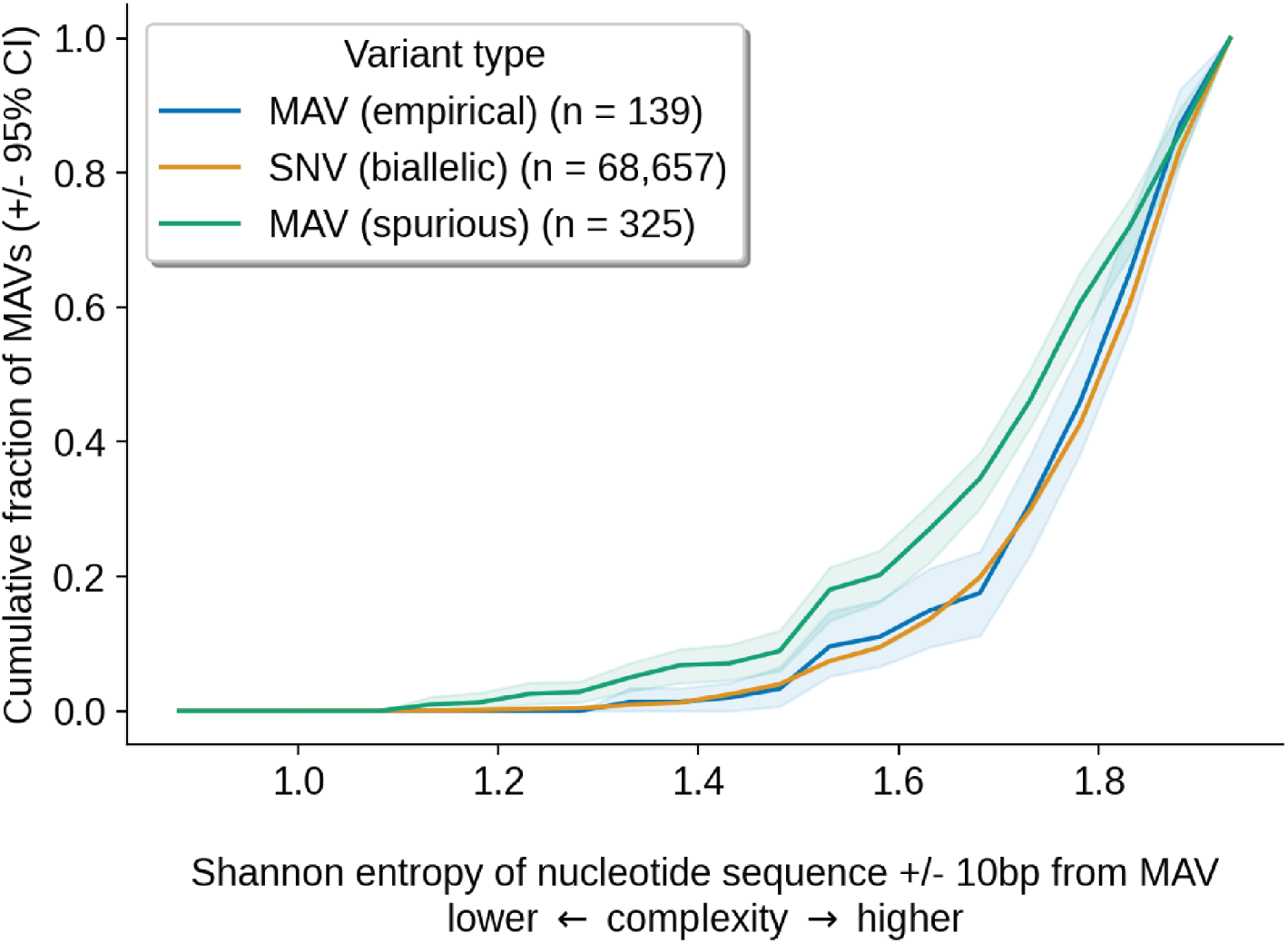
Empirical MAVs occur in higher-complexity sequences than spurious MAVs. We used `mutyper` [46] to extract the flanking 10 nucleotides of sequence context surrounding every MAV with at least three reads of support (including *n* = 139 “empirical” MAVs and *n* = 325 “spurious” MAVs identified using a random strain’s sequencing reads). We then calculated the Shannon entropy of the 20bp nucleotide context (excluding the mutated nucleotide) at each MAV. We also calculated the entropy of the 20bp nucleotide context surrounding all *n* = 68,657 biallelic SNVs (excluding MAVs) observed in DMS, EMS, and MMS-treated strains.

**Supplementary Figure 3:**
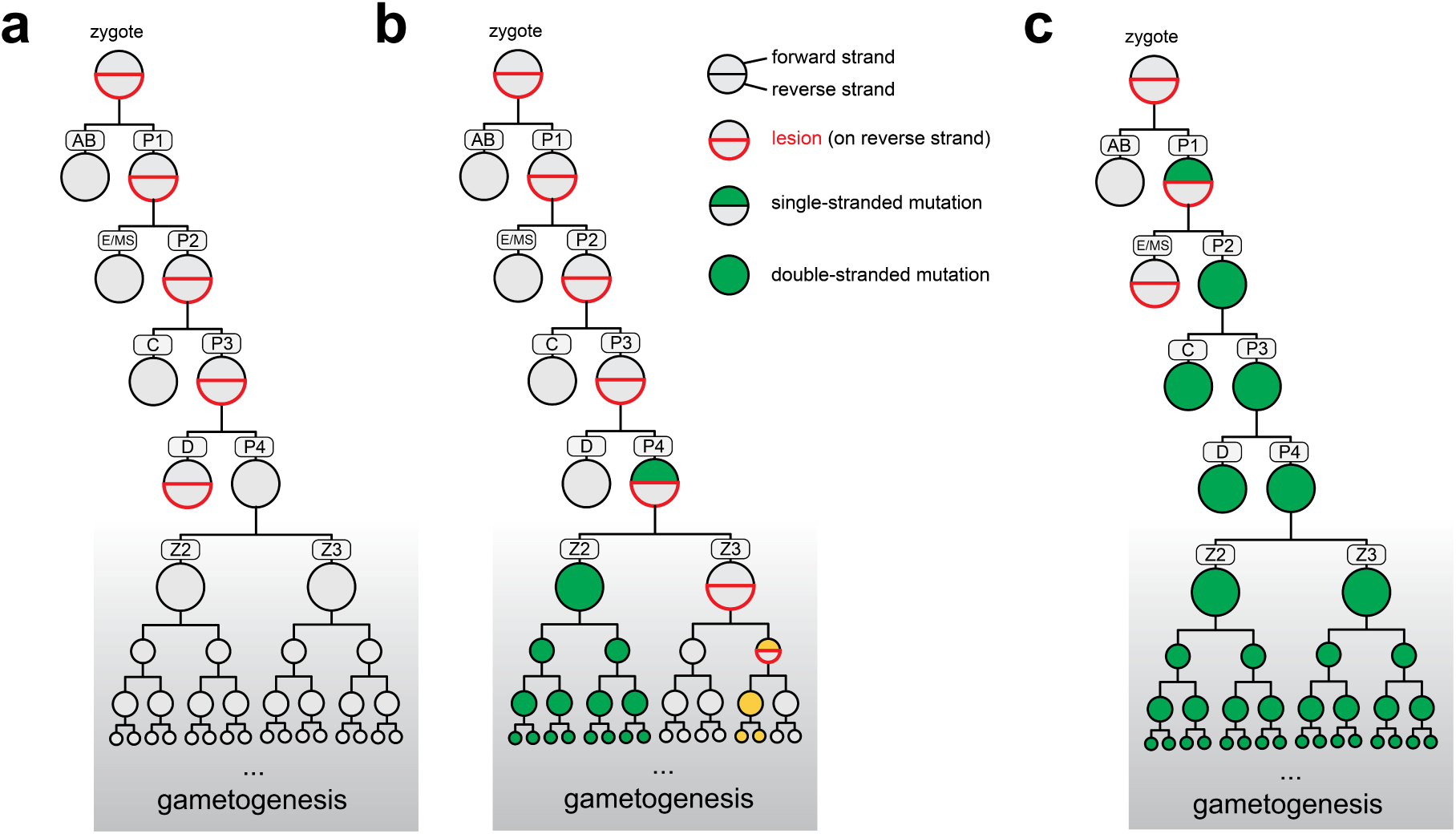
Schematic of lesion segregation during *C. elegans* embryogenesis. In each panel, we depict the fate of a single nucleotide on the paternally-inherited chromosome in an F1 embryo (the offspring of a mutagenized P0). These illustrations are simplified versions of the schematic shown in Figure 5. We do not depict the maternally-inherited chromosome, which is assumed to be lesion-free. Single-nucleotide sites are depicted as half-circles; the top half corresponds to the forward strand and the bottom half corresponds to the reverse strand. Half-circles are outlined in red if that strand carries a lesion at that site. Half-circles are colored grey if they possess a “reference” (non-mutated) nucleotide at the site, and either green or gold if they possess a non-reference nucleotide. Circles are fully colored if a mutation is “fully-resolved” (i.e., double-stranded). **a)** If a lesion-containing strand segregates into a somatic progenitor cell (or undergoes error-free repair) prior to the 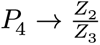 cell division, no gametes will possess a mutation. **b)** If the lesion templates an incorrect nucleotide (colored in green) during the *P*_3_ → *P*_4_ division, segregates unrepaired into gamete progenitors, and later templates the incorporation of a second incorrect nucleotide, the second *de novo* mutation (colored in gold) will be present in a comparatively small fraction of mature gametes. **c)** If the lesion templates an incorrect nucleotide during an early embryonic cell division, and the incorrect nucleotide continues to segregate into the gamete progenitors (i.e., the *P* lineage), half of all mature haploid gametes will possess the mutation.

**Supplementary Figure 4:**
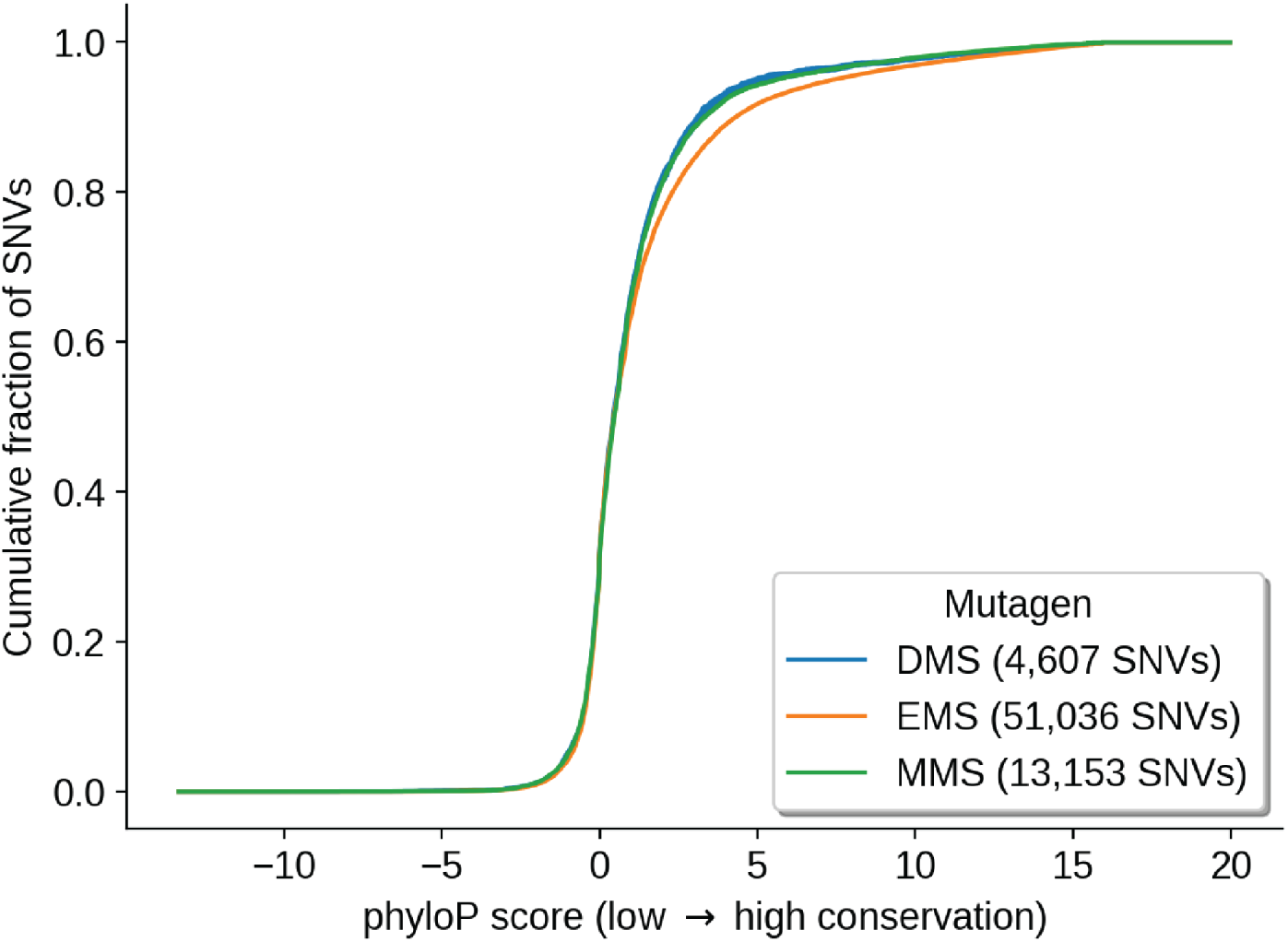
EMS-derived mutations occur in more conserved regions of the genome. We measured the phyloP [47] score (using the `phyloP135way` track for the `ce11` reference assembly, downloaded from the UCSC Table Browser) at every SNV identified in strains treated with DMS, EMS, and MMS in Volkova *et al.* (2020). Here, we show the cumulative fraction of SNVs with a phyloP score of at least *x*. Two-sided Kolmogorov-Smirnov *p*-values from comparisons of phyloP score distributions between DMS and EMS: 2.3 × 10^−13^; between DMS and MMS: 0.3; between MMS and EMS: 1.9 × 10^−28^.

